# GNMCADS: Sampling For Conformation Diversity With Gaussian Network Model Guided Condition Annealed Diffusion Sampler

**DOI:** 10.64898/2026.08.28.747885

**Authors:** Ahmed Selim Uzum, Turkan Haliloglu

## Abstract

Biomolecules exist in diverse conformations underlying their biological functions. Although recent approaches on molecular dynamics emulation, evolutionary information perturbation, or diffusion steering enabled sampling of distinct conformations, capturing major domain motions or long-timescale transitions still remains a challenge. Here, we introduce GNMCADS, an inference-time conformational sampling strategy that enhances the sampling diversity of biomolecular diffusion models. Using the intrinsic dynamics of biomolecules, GNMCADS identifies dynamical domains of biomolecular systems, revealing residues with correlated and anti-correlated motions, and selectively anneals the diffusion conditioning signal, informed by these dynamical domains allowing the sampling of diverse yet biologically relevant conformations. When implemented in AlphaFold3, GNMCADS exhibits improved conformational coverage compared to other conformational sampling methods, without requiring additional training or sampling with multiple seeds, across a set of 92 proteins comprising GPCRs, transporters, and proteins undergoing large domain movements . Similarly, GNMCADS captures experimentally resolved conformations that AlphaFold3 cannot sample for the de novo designed rhodamine binder, which lacks tractable evolutionary information. Beyond proteins, GNMCADS also uncovers experimentally resolved HIV-1 TAR and RNase P RNA conformations that cannot be sampled by AlphaFold3. Together, these results highlight the potential of utilizing intrinsic dynamics during diffusion to sample diverse biologically relevant conformations without additional training.

## 1 Introduction

Proteins are the essential machines that regulate and execute cellular activities. With the recent advent of deep learning methods such as AlphaFold2 [1], and its successor AlphaFold3 (AF3) [2], it has become possible to generate the three-dimensional structure of a protein from its sequence. However, proteins are inherently dynamic molecules adopting multiple conformational states which dictate their biological functions. Hence, the characterization of the conformational landscapes of proteins is crucial for understanding their biological functions [3].

Recent machine-learning based approaches aiming to tackle the problem of conformational sampling from sequence and evolutionary information alone can be classified into three main categories. The first category includes methods trained or fine-tuned using molecular dynamics (MD) simulations to learn to sample protein conformations from their underlying free energy landscapes [4–7]. However, the prohibitive cost of MD simulations limits the availability of training data that capture conformational changes occurring in large protein complexes or over long timescales. Consequently, these methods may have difficulty sampling biologically relevant conformational changes that are underrepresented in their training data. In contrast to methods in the first category, methods in the second and third categories do not require training with MD simulations and increase the conformational sampling diversity of already trained protein structure prediction models such as AlphaFold. Methods in the second category accomplish this by modifying the multiple sequence alignments (MSAs) used as the source of coevolutionary information by structure prediction models. Specifically, they generate an ensemble of structures by altering the coevolutionary information provided to the structure prediction models by subsampling MSAs while varying the number of aligned sequences [8], clustering aligned sequences by their sequence similarity, and generating new MSAs from these clusters [9], or performing column-wise MSA modifications [10–12]. Whereas the second category increases diversity by manipulating the inputs of the prediction models, the third category achieves diversity by internal interventions in the prediction models by steering latent representations [13] or guiding the diffusion sampling process [14]. Compared to increasing sampling diversity by MSA perturbations, operating on internal model representations enables these methods to avoid relying directly on the availability of evolutionary information in terms of MSA depth or heterogeneity which is advantageous for proteins with low evolutionary information such as de novo designed proteins.

In this study, we introduce GNMCADS, a diffusion sampler that enhances the conformational sampling diversity of biomolecular diffusion models by annealing pairwise conditioning signal in a selective manner based on the intrinsic dynamical organization of sampled structures (Fig. 1a). Although condition annealing has been successfully used to increase the sampling diversity of image diffusion models [15], naively implementing condition annealing can have substantial effects on biomolecular diffusion models, resulting in profound decrease in sampling accuracy (Fig.2c, 2d). To this end, GNMCADS anneals the diffusion conditioning signal selectively by taking the intrinsic dynamical properties of the sampled conformations into account. Specifically, this is achieved by generating a reference structure using default sampling, and leveraging the Gaussian Network Model’s (GNM) ability to predict the equilibrium fluctuations and identify large-scale biologically functional collective motions that are intrinsically embedded in the sampled reference structure [16–19]. Subsequently, GNMCADS divides the initially predicted reference structure into distinct ”dynamical domains” based on the cross-correlations of residue fluctuations inferred from GNM such that the residues in a dynamical domain tend to exhibit correlated motions whereas the residues in different dynamical domains tend to exhibit anti-correlated movements. Finally, the calculated dynamical domains are used to guide the annealing process with the addition of noise specifically to the inter-dynamical domain and non-diagonal elements of the pairwise conditioning signal, enabling the denoising process to generate conformations that are compatible with the underlying intrinsic dynamics of the sampled biomolecule.

**Figure 1:**
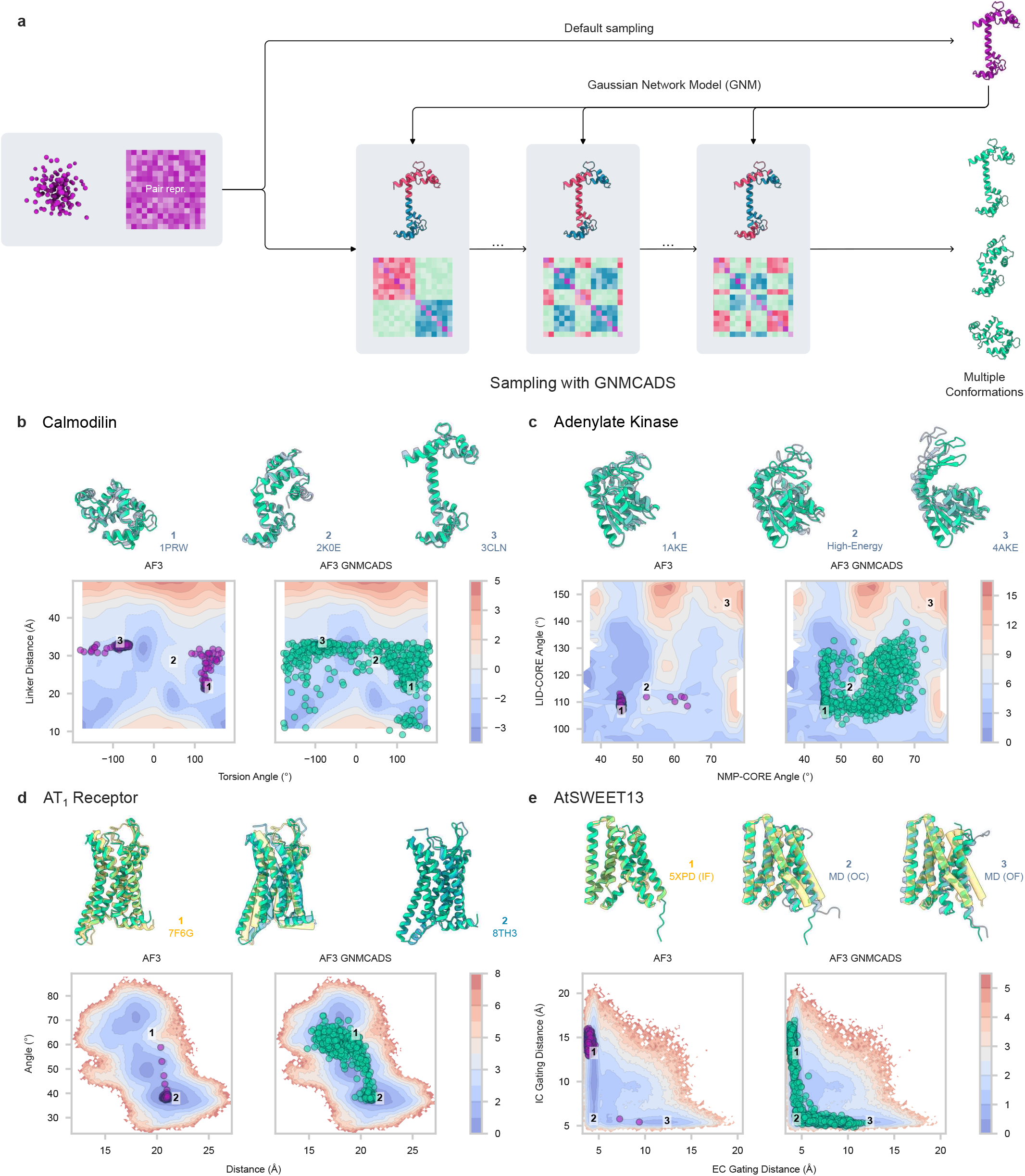
GNMCADS enables sampling of diverse protein conformations in diffusion models. **a** GNMCADS samples diverse protein conformations by annealing the pairwise conditioning signal between the intrinsic dynamical domains of the initially sampled structure identified by GNM. The opposing dynamical domains are represented by red and blue, the annealed section of the pairwise representation is shown in light green. **b-e** Structures generated using default AF3 (purple) and AF3 GNMCADS (green), projected on their corresponding free energy landscapes and superimposed onto the experimentally resolved or MD derived conformations (grey). The landscape color scales represent the relative free energy in kcal/mol. See Section S1.1 for further information about the superimposed structures and the free energy landscapes used.

Further, we present AF3 GNMCADS, an implementation of GNMCADS in the diffusion module of AlphaFold3 and benchmark its conformational sampling capabilities over three datasets that consist of 92 proteins across 54 class A GPCRs, 15 transporters and 23 proteins with major conformational changes. We compare AF3 GNMCADS sampling against AlphaFold3’s default sampling; Boltz2 in its MD emulation mode [6] and BioEmu [4], both of which are methods trained on MD simulations; AFSample2 [11] and AFSample3 [12], which perform column-wise MSA randomizations in AlphaFold2 and AlphaFold3 respectively; and ConforMix [14] which employs RMSD-based bias potentials during the diffusion process. Moreover, we demonstrate that GNMCADS can enhance the sampling of not only naturally occurring proteins but also de novo designed proteins and RNA systems, capturing their experimentally observed conformations that AlphaFold3’s default sampling strategy is unable to predict.

## 2 Results

### 2.1 GNMCADS captures biologically relevant conformations

For increased diversity in biomolecular generative models to be useful, the generated structures must represent distinct biologically relevant conformations of the sampled protein. We assess GNMCADS’ ability to generate such conformations by projecting the generated structures of calmodulin, adenylate kinase, AT1aR GPCR, and AtSWEET13 transporter onto their respective MD simulation derived free energy landscapes parameterized by the collective variables that are crucial for their biological functions (Fig. 1b-e) [20–23]. Across all four proteins, AF3 GNMCADS samples a broader range of biologically relevant conformational states and more thoroughly populates their respective free energy landscapes compared to AF3.

Notably, AF3 GNMCADS captures extended, compact, and intermediate conformations of calmodulin; closed, open, and PCS–CPMG NMR-derived [24] high energy conformations of adenylate kinase; open, intermediate, and closed conformations of AT1aR GPCR; and inward-facing, occluded, and outward-facing conformations of AtSWEET13 transporter. Moreover, AF3 GNMCADS sampled structures populate regions of the free energy landscapes that span between these conformational states, indicating that AF3 GNMCADS also captures intermediate conformations along functionally relevant transitions.

### 2.2 Benchmarking AF3 GNMCADS against other conformational sampling methods

To assess the necessity of utilizing inter-dynamic domain masking and further evaluate GNMCADS’ conformational sampling abilities, we benchmarked GNMCADS on the open-closed [11] and transporters [25] datasets which comprise 23 proteins with distinct open and closed conformations, and 15 transporters with inward-facing and outward-facing conformations, respectively, showcasing large domain motions (Fig. 2).

**Figure 2:**
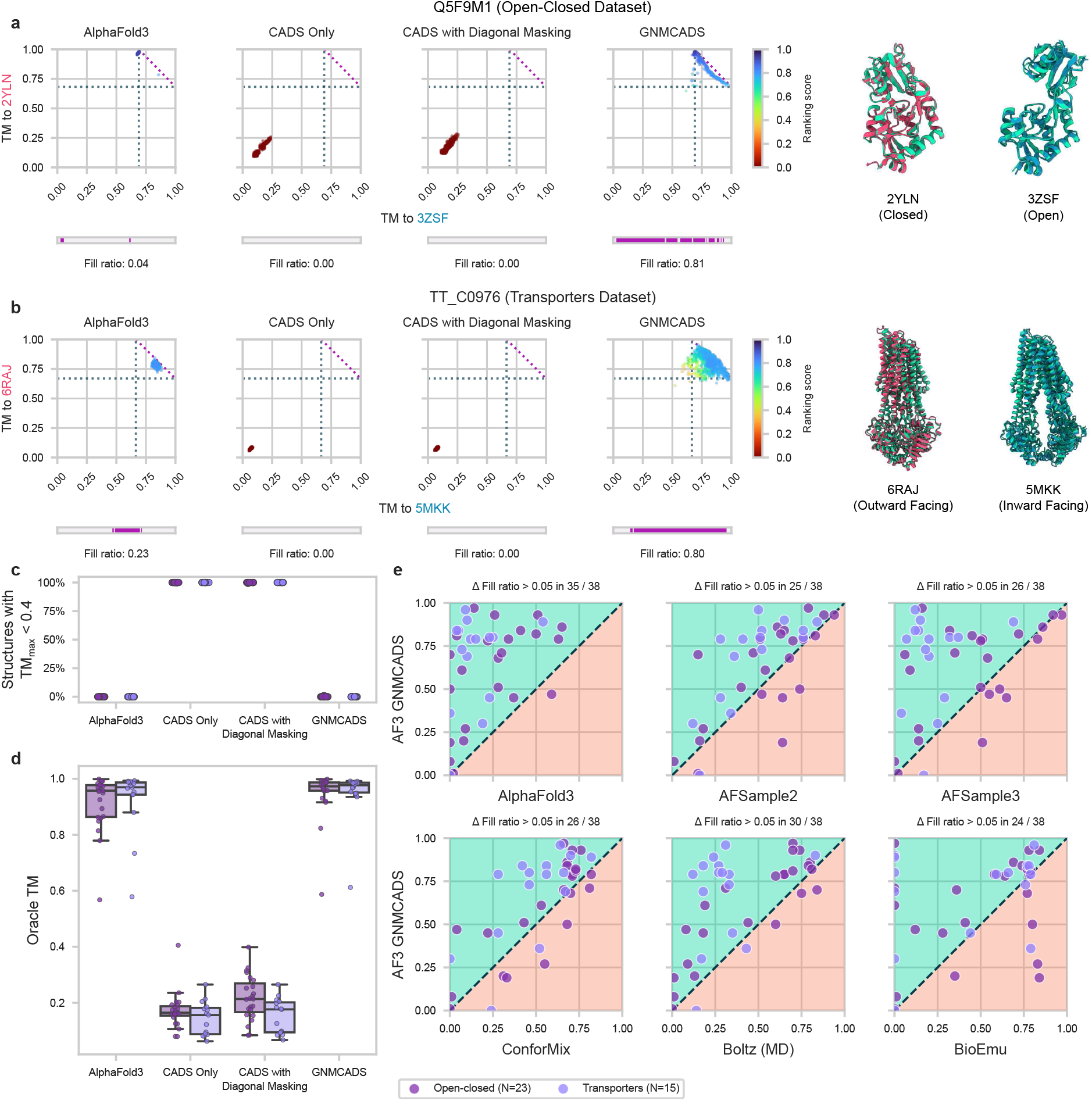
GNMCADS increases sampling diversity compared to other methods across varying targets. **a**, **b** Representative cases Q5F9M1 and TT_C0976 for the open-closed and transporters datasets, respectively. The left column shows the TM similarity and fill ratio plots for structures predicted by the default AF3 sampling (AF3), AF3 with CADS sampling (CADS Only), AF3 with CADS sampling with diagonal masking (CADS with Diagonal Masking) and AF3 GNMCADS. The right columns show AF3 GNMCADS predicted structures (green) superimposed onto the experimentally determined conformations. **c** Percentage of the predicted structures with lower than 0.4 TM score to any experimentally resolved conformations. **d** Oracle TM scores for each target, which is defined as the highest TM score between any predicted and any experimental structure. **e** The per-target fill ratios of conformational sampling methods compared against AF3 GNMCADS. Δ Fill ratio *>* 0.05 represents the number of targets for which AF3 GNMCADS improves their fill ratio by more than 0.05.

**Figure 3:**
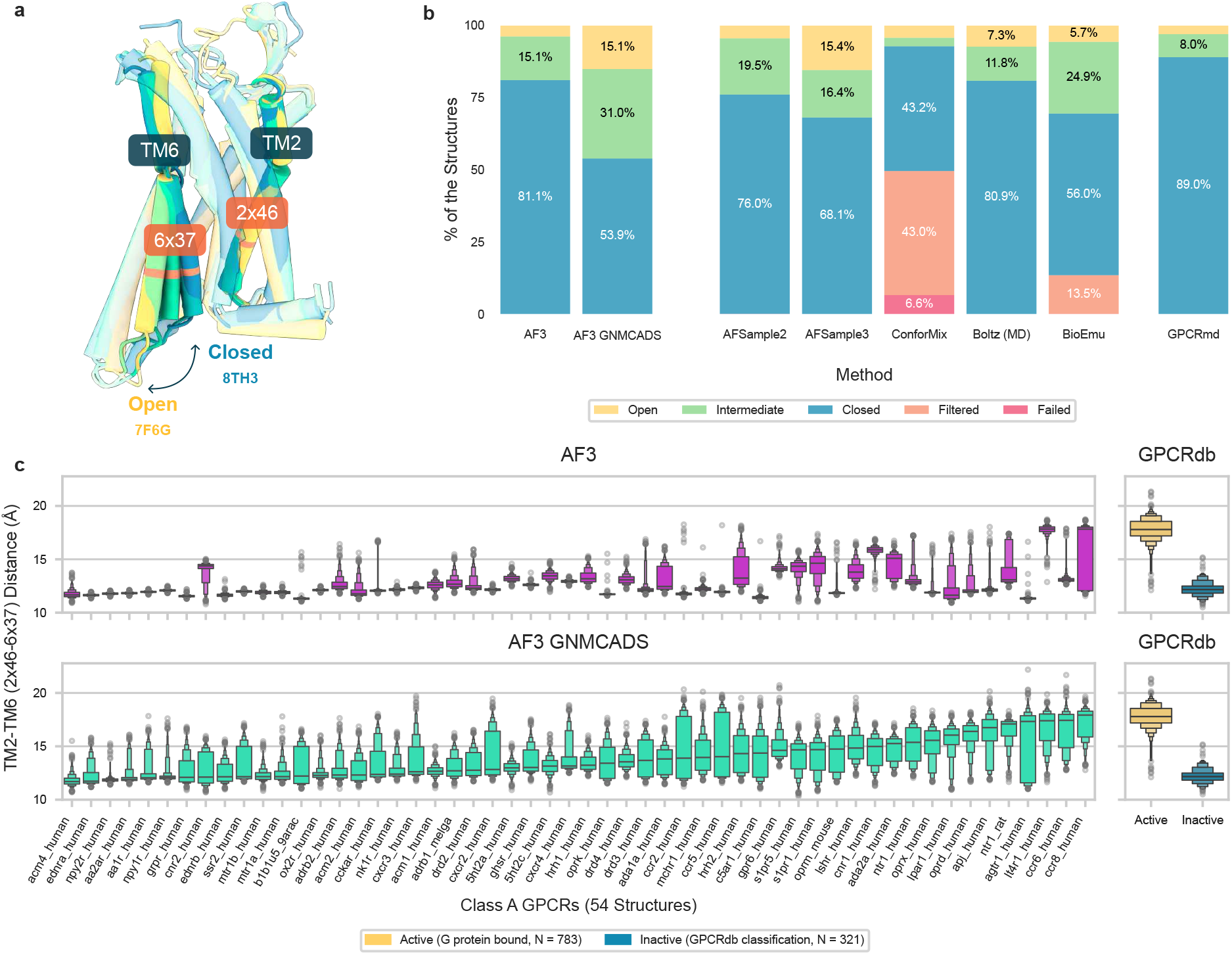
AF3 GNMCADS samples diverse GPCR conformations. **a** Open (7F6G, [28]) and closed (8TH3, [29]) conformations of AT1aR, superimposed onto AF3 GNMCADS sampled open, intermediate, and closed conformations (green). **b** Percentage of conformations generated by AF3, AF3 GNMCADS and other conformational sampling methods across 54 class A GPCRs, and 3 × 500ns of MD simulations of 30 out of the 54 GPCRs performed by GPCRmd [26, 30]. For clarity, percentages less than 5% are not shown. See Table S3 for further details. **c.** TM2-TM6 distances of the structures sampled by AF3 and AF3 GNMCADS, compared with the TM2-TM6 distances of experimentally determined GPCR structures classified by GPCRdb [27] as G protein bound or inactive. The TM2-TM6 distances are shown as letter-value plots [31], in which the central line represents the median, the innermost box spans the first to third quantiles and the outer boxes represent the successive quantiles.

On both of the tested datasets, removing GNM derived inter-dynamic domain masks resulted in substantial degradation of the accuracy of predicted proteins for which nearly 100% of the structures sampled via CADS or CADS with diagonal masking had a TM score of less than 0.4 against every experimentally determined conformation for a target (Fig. 2c) Furthermore, we used the fill ratio [11] to quantitatively compare the conformational sampling ability of AF3 GNMCADS and other conformational sampling methods. Across 38 proteins benchmarked among both datasets, AF3 GNMCADS improved the fill ratio by more than 0.05 for the majority of the targets across all other tested conformational sampling methods.

### 2.3 AF3 GNMCADS recovers diverse conformations along GPCR activation

We evaluated the ability of AF3 GNMCADS to capture long-timescale conformational transitions by benchmarking it against AF3 and other conformational sampling methods across 54 class A GPCRs, as the conformational state transitions of GPCRs occur on multi-microsecond timescales [26]. Subsequently, we categorized the conformational states of the sampled structures according to their TM2 to TM6 helix distance, a collective variable that represents the opening of the internal cavity during activation of a GPCR [26, 27] (see Section 4.4).

Across the 54,000 GPCR structures sampled with each method, AF3 GNMCADS substantially broadens per-GPCR TM2-TM6 distance ranges by generating an increased number of structures in open and intermediate conformations than AF3, which predominantly generates GPCR structures in closed conformations. Specifically, 81.1% of the AF3 sampled GPCR structures were classified as closed, whereas only 15.1% and 3.8% were classified as intermediate and open conformations, respectively. In contrast, AF3 GNMCADS captures considerably more diverse GPCR conformations than AF3, with 53.9%, 31.0%, and 15.1% of the sampled structures being classified as closed, intermediate, and open conformational states, respectively. Furthermore, when compared against other benchmarked conformational sampling methods, AF3 GNMCADS also samples the largest number of GPCRs in their intermediate conformations.

### 2.4 AF3 GNMCADS enhances conformational sampling of RNA structures and de novo designed proteins

Next, we sought to test GNMCADS on biomolecular systems beyond naturally existing proteins by assessing its conformational sampling abilities on de novo designed proteins and RNA structures (Fig. 4). Compared to naturally existing proteins, de novo designed proteins do not have evolutionary information that can be tractable with multiple sequence alignments. Hence, many of the conformational sampling methods that are solely based on perturbing multiple sequence alignments are unable to effectively sample their conformational landscapes. To this end, we tested AF3 GNMCADS on Rhobin9, a de novo designed rhodamine binder, which has merely one aligned sequence in its multiple sequence alignment (Fig. 4a). When compared to AlphaFold3’s default sampling, which was only able to sample the holo form of the Rhobin9, GNMCADS was able to recover both the apo and holo forms in combination with intermediate forms as well.

**Figure 4:**
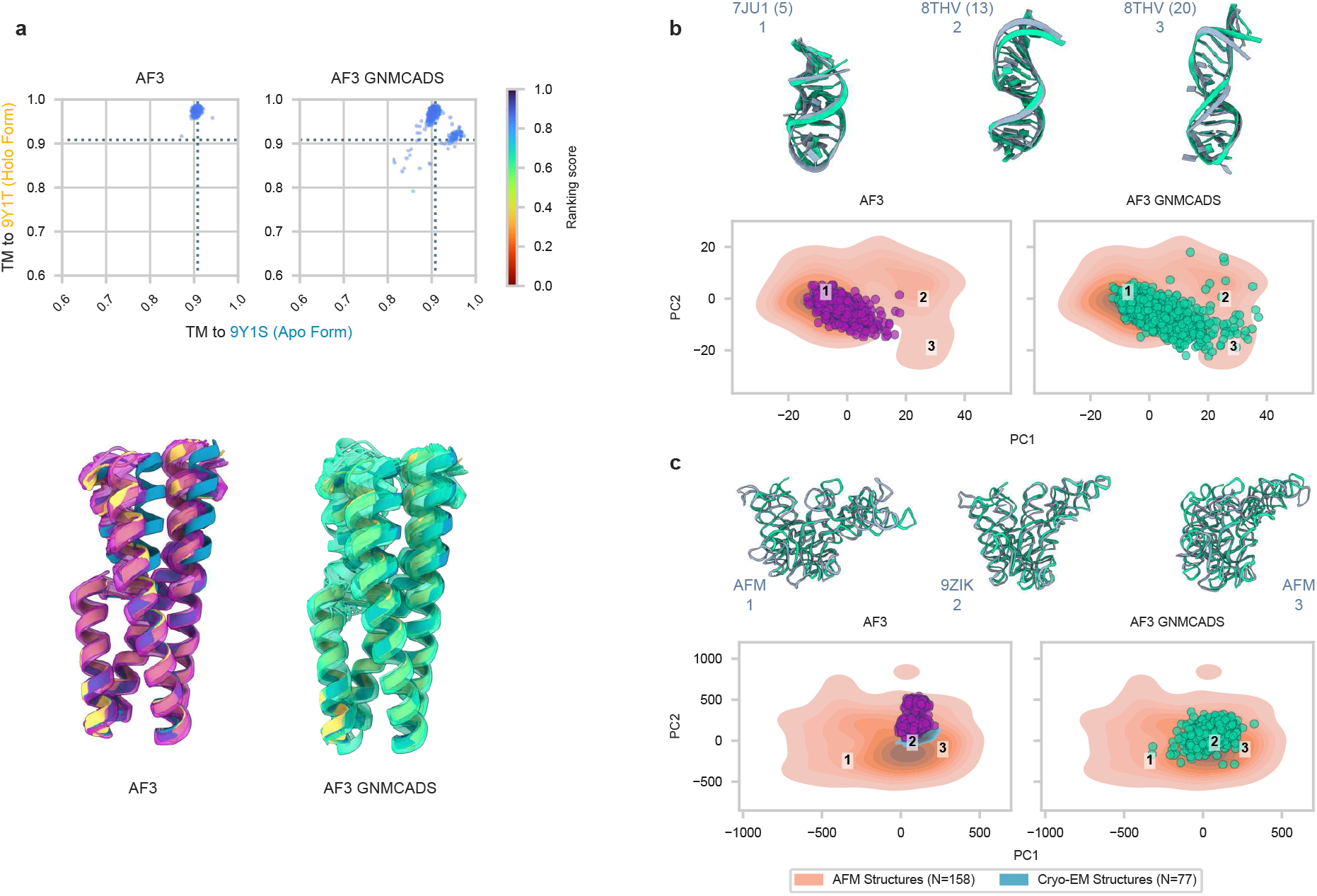
AF3 GNMCADS enhances RNA and de novo designed protein conformation sampling. **a** Comparison of AF3 and AF3 GNMCADS sampled structures for the de novo designed rhodamine binder, Rhobin9. The AF3 sampled structures are shown in purple, AF3 GNMCADS sampled structures are shown in green. The experimentally resolved apo and holo Rhobin9 structures are shown in blue and yellow, respectively. **b,c** Visualisation of GNMCADS sampled HIV-1 TAR and RNase P conformations, respectively via principal component analysis (PCA). AF3 sampled structures are represented in purple, AF3 GNMCADS sampled structures are represented in green, and experimentally determined structures are represented in grey. See Section S2.1 for further information about principal component analysis.

Following the previous works utilizing Gaussian Network Models for analysing dynamics of RNA structures [32, 33], we also tested GNMCADS’ ability to sample diverse RNA conformations on HIV-1 TAR and RNase P structures (Fig. 4b, Fig. 4c). Similar to other tested proteins, sampling with GNMCADS was able to capture distinct experimentally resolved conformations of both HIV-1 TAR and RNase P compared to AlphaFold3, better exploring their respective principal component conformational landscapes.

## 3 Discussion

Structure prediction models can accurately generate structures of biomolecules such as proteins or nucleic acids from their sequences, yet they remain limited in capturing their conformational variability. GNMCADS addresses this limitation by incorporating a physically motivated description of intrinsic collective dynamics into the diffusion sampling process by selectively relaxing pairwise constraints between GNM-identified dynamical domains while preserving internal coupling via condition annealing. This targeted strategy enhances sampling along collective structural rearrangements without requiring additional training.

When evaluated against molecular dynamics (MD) derived free energy landscapes of calmodulin, adenylate kinase, AT1aR GPCR, and AtSWEET13 transporter, AF3 GNMCADS, the AlphaFold3 implementation of GNMCADS, achieves substantially broader coverage along the functionally relevant regions of their respective free energy landscapes and generates a greater number of distinct biologically relevant conformational states compared to AlphaFold3. The increased sampling of intermediate states is particularly notable, as biologically relevant conformational transitions often involve coordinated rearrangements between partially populated states. The GPCR analysis further illustrates this effect: AF3 strongly favored closed conformations, whereas sampling with GNMCADS substantially increased the representation of intermediate and open states and sampled intermediate conformations across more receptors. In addition, as GNMCADS employs intrinsic dynamics of proteins to sample their conformational landscape, it does not depend on the availability of evolutionary information, allowing the sampling of orphan proteins such as de novo designed proteins. Similarly, when applied to RNA structures HIV-1 TAR and RNase P, sampling with GNMCADS enables the prediction of multiple experimentally observed conformational states that are not sampled via AlphaFold3.

Moreover, when benchmarked across 92 targets including class A GPCRs, transporters, and proteins that undergo major domain movements, AF3 GNMCADS achieves state-of-the-art conformational sampling performance compared to other conformational sampling methods based on multiple sequence alignment (MSA) modification, MD emulation, and twisted diffusion sampling. The consistent improvement across these distinct approaches suggests the importance of incorporation of dynamic organization to diffusion sampling via condition annealing.

Together, these results demonstrate how integrating intrinsic biomolecular dynamics with condition annealing into diffusion models can expand the exploration of conformational ensembles of proteins and nucleic acids, providing a physics-aware route toward understanding the structural diversity underlying biomolecular function.

## 4 Methods

### 4.1 Determination of dynamical domains with GNM

The Gaussian Network Model (GNM) is a minimalist physics-based coarse-grained normal mode model that describes the equilibrium dynamics of proteins and nucleic acids [16, 17, 32, 33].

In GNM, biological systems consisting protein and nucleic acids are represented as an elastic network in which C*_α_* atoms of proteins and phosphorus atoms of nucleic acids that lie within a cutoff distance, *r_c_*, are connected by uniform harmonic springs. Under the assumption of Gaussian-distributed fluctuations, this representation enables the characterization of equilibrium motions within a statistical thermodynamics framework, where collective residue fluctuations are decomposed into intrinsic normal modes of the network. This formulation enables the equilibrium cross-correlations of fluctuations between residues *i* and *j* to be expressed as:

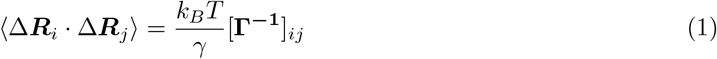

where ***R****_i_*is the position vector of the *i*th residue’s representative atom, *k_B_*is the Boltzmann constant, *T* is the absolute temperature, *γ* is the force constant, and **Γ** is the Kirchhoff matrix representing the connectivity of the protein such that

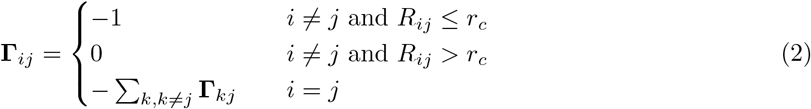

where *R_ij_*is the distance between the representative atoms of the residues *i* and *j*.

The pseudo-inverse of **Γ** can be represented by the sum of the *N* − 1 non-zero modes 2 ≤ *k* ≤ *N* using

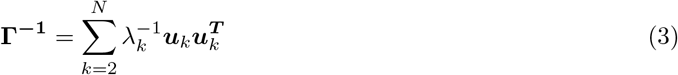

where *λ_k_* and ***u****_k_* are the eigenpairs of **Γ**. The first mode *k* = 1 is the trivial mode with *λ*_1_ = 0 and is omitted.

Since ⟨Δ***R****_i_* · Δ***R****_j_*⟩ is directly proportional to 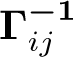, the cross-correlation of the equilibrium fluctuations between the residues *i* and *j* for the mode *k* can be calculated as:

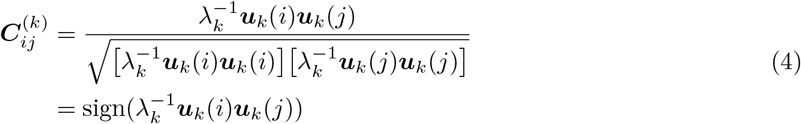

Using the per mode cross-correlations, a given biological system consisting of protein or nucleic acids can be divided into two dynamical domains where the residue pairs within the same dynamical domain tend to exhibit correlated motions and have positive cross-correlation values, whereas the residue pairs belonging to the opposing dynamical domains tend to exhibit anti-correlated motions and have negative cross-correlation values.

### 4.2 GNMCADS

GNMCADS employs two masks to perform selective annealing. The first mask, ***M***_diag_, is used to limit the annealing of the diagonal entries along with the first *k*_diag_th superdiagonal and subdiagonal entries of the pairwise conditioning signal as it has been shown that the perturbations of the diagonal entries of the pairwise conditioning signal can have substantial effects on the secondary structure of the generated proteins [34]. The ***M***_diag_ mask is defined as:

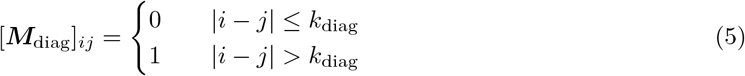

The second mask used by GNMCADS, ***M***_GNM_, enables the annealing process to become aware of the intrinsic dynamics of the sampled biomolecule. This is achieved by performing the annealing process only for the entries of the pairwise conditioning signal that contain inter dynamical domain information. As the residues in the opposing dynamical domains tend to have anti-correlated motions, annealing of the entries that belong to the inter dynamical domain region of the pairwise conditioning signal allows GNMCADS to sample diverse conformations that are compatible with the sampled biomolecule’s underlying intrinsic dynamical behaviours. The ***M***_GNM_ mask for the GNM mode *k* is defined as:

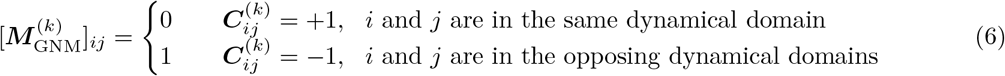

These two masks are then combined as a single mask 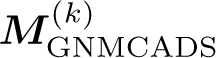 such that:

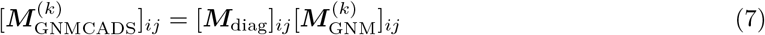

Low-frequency GNM modes encode the collective functional motions and many experimentally observed conformational transitions can thus be well described by the lowest frequency GNM modes [35–37]. Accordingly, for each diffusion step *t*, the value of *k_t_*, which is the GNM mode that will be used to define dynamical domains and consequently the mask that will be used is selected from a distribution that favors low-frequency modes such that:

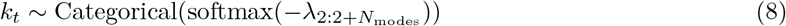

Subsequently, the newly selected GNM mode *k_t_* is then used to anneal the pairwise conditioning signal ***z***^trunk^ ∈ R*^N×N×Nc^* where *N_c_* is the channel dimension of the pairwise conditioning signal. However, unlike CADS, GNMCADS also scales the noise values ***n****_ijc_*∼N (0, 1) with the mean and standard deviation of each channel’s selected elements. The complete annealing step of GNMCADS can be shown as:

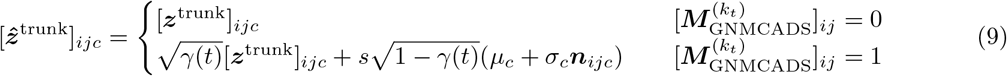

where *s* is the noise scale, *µ_c_*and *σ_c_* are the mean and the standard deviation of the elements selected by 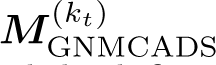 in the *c*^th^ channel of the pairwise conditioning signal, and *γ*(*t*) is the linear annealing schedule defined as:

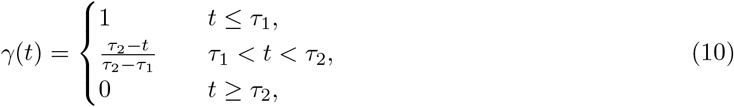

where the values of *τ*_1_*, τ*_2_ ∈ (0, 1) are the thresholds of the linear annealing schedule.

### 4.3 Inference protocols

Each benchmarked method was used to generate 1000 structures for each target. AF3 predicted structures were generated by rerunning the model with 200 seeds and producing 5 diffusion samples per seed. In contrast, AF3 GNMCADS predicted structures were generated using one seed with 1000 diffusion samples per seed.

AFSample2, AFSample3, and BioEmu were used as described in their original studies. For ConforMix, 1000 structures were generated instead of the 800 structures recommended by its authors. Similarly, the Boltz2 (MD) method was used to generate 1000 structures by using 200 seeds with 5 diffusion samples per seed.

### 4.4 GPCR state determination

Conformational states of the predicted GPCR structures were determined using the TM2-TM6 distance thresholds proposed by [26] which are calculated as the distances between the 2×46 and 6×37 residues in the GPCRdb numbering scheme [27]. The TM2-TM6 distance thresholds of 13.6 Å and 16.36 Å were used for the intermediate and open conformations, respectively; structures with TM2-TM6 distances less than 13.6 Å are classified as closed structures.

Predicted structures with a 7TM RMSD (excluding the extracellular and intracellular loops) greater than 7 Å relative to both the G protein bound and inactive experimentally determined structures were classified as failed predictions.

### 4.5 Fill ratio

The fill ratio metric, first introduced in [11], assesses the protein generative models based on their ability to sample contiguous transitions between two known conformations of a protein using the TM similarity scores [38].

To calculate the fill ratio for a target with experimentally resolved conformations *X* and *Y* , the TM scores between predicted structures and the reference structures *X* and *Y* are calculated with US-align [39], using a length-dependent *d*_0_ normalization factor selected by US-align based on the sequence length of the reference structure to mitigate the dependency between sequence lengths and the similarity scores [38]. Using the calculated TM scores, the structures including the reference structures, are represented as points in two dimensions with coordinates (TM score to structure *X*, TM score to structure *Y*). The predicted structures having greater TM similarity score to both structures *X* and *Y* compared to the TM similarity score between *X* and *Y* structures themselves are labeled as intermediate structures and projected onto the diagonal that connects the points representing the TM scores between the experimental conformations *X* and *Y* . Finally, the histogram of the projected structures is calculated using 100 equal width bins and the fraction of the bins that have at least one structure is calculated as the fill ratio of the target.

## Code availability

The source code for GNMCADS, and its AlphaFold3 implementation AF3 GNMCADS is available at https://github.com/ahmedselim2017/af3_gnmcads.

## Acknowledgments

We would like to thank Sergey Ovchinnikov and Ezgi Karaca for their valuable discussions.

The numerical calculations reported in this paper were partially performed using the EuroHPC Joint Undertaking (EuroHPC JU) supercomputer MareNostrum 5, hosted by the Barcelona Supercomputing Center (BSC). Access to MareNostrum 5 was provided through a national access call coordinated by the Scientific and Technological Research Council of Turkiye (TUBITAK). We gratefully acknowledge BSC, TUBITAK, and the EuroHPC JU for providing access to these resources and supporting this research. This work was supported by the Bogazici University BAP Research Fund 25A05P4.

## Author contributions

Conceptualization: A.S.Ü., T.H.; Methodology: A.S.Ü., T.H.; Software: A.S.Ü.; Validation: A.S.Ü., T.H.; Data Curation: A.S.Ü.; Writing: A.S.Ü., T.H.; Supervision: T.H.; Project administration: T.H.; Funding acquisition: T.H.

## Supplementary Information

### S1 Datasets

#### S1.1 Free energy landscapes

The free energy landscapes used in this study and their references are shown in Table S1.

**Table S1:** References for the free energy landscapes used.

| Protein | Reference |
|---|---|
| Calmodulin | [1] |
| Adenylate kinase | [2] |
| AT1aR | [3] |
| AtSWEET13 | [4] |

The experimentally resolved structures compared with AF3 GNMCADS predicted structures and projected onto their respective free energy landscapes are shown in Table S2.

**Table S2:**
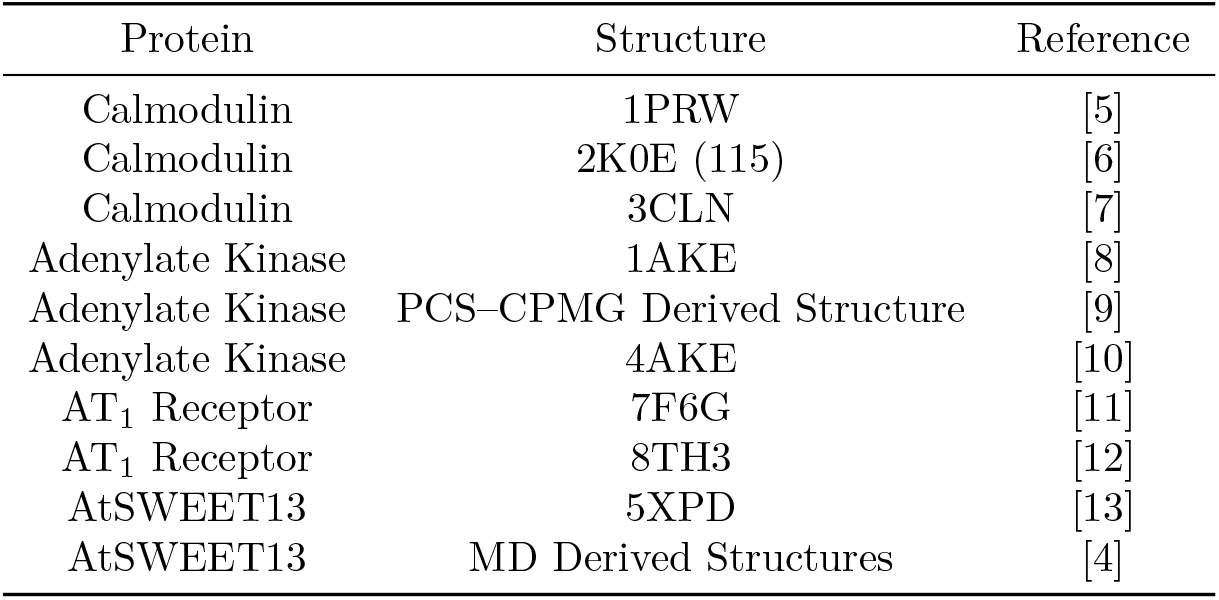
References for the structures projected to their respective free energy landscapes.

| Protein | Structure | Reference |
|---|---|---|
| Calmodulin | 1PRW | [5] |
| Calmodulin | 2K0E (115) | [6] |
| Calmodulin | 3CLN | [7] |
| Adenylate Kinase | 1AKE | [8] |
| Adenylate Kinase | PCS–CPMG Derived Structure | [9] |
| Adenylate Kinase | 4AKE | [10] |
| AT<sub>1</sub> Receptor | 7F6G | [11] |
| AT<sub>1</sub> Receptor | 8TH3 | [12] |
| AtSWEET13 | 5XPD | [13] |
| AtSWEET13 | MD Derived Structures | [4] |

#### S1.2 GPCR dataset

The GPCR dataset was created by applying the following filters to the GPCRdb database [14]:

- The GPCR must be a class A GPCR.
- Both the G protein bound and inactive states of the GPCR must be available.
- The inactive state must have a GPCRdb defined activation degree of 10% or less.
- The GPCR structures must have a resolution of 9 Å or better.

For GPCRs with multiple active or inactive states after the filtering, the structure with the highest sequence similarity to the canonical sequence was selected. Structure predictions were performed using the canonical 7TM (TM1-H8) sequences including the extracellular and the intracellular loops defined by GPCRdb.

#### S1.3 Open-closed dataset

The open-closed dataset introduced by [15] contains proteins that undergo large-scale domain motions. As the target Q53W80 exists in both monomer and homodimer conformations, its homodimer form has also been included in the dataset. Similarly, as the target B7IE18 is also included in the transporters dataset, it has been removed from the open-closed dataset. The PDB identifiers for the targets in the dataset can be seen in Table S4.

**Table S3:** Conformational states generated across 1000 predicted structures over 54 class A GPCRs.

| Method | Open | Intermediate | Closed | Filtered | Failed |
|---|---|---|---|---|---|
| AF3 | 2,045 | 8,177 | 43,778 | 0 | 0 |
| AF3 GNMCADS | 8,152 | 16,738 | 29,109 | 0 | 1 |
| AFSample2 | 2,402 | 10,524 | 41,059 | 0 | 15 |
| AFSample3 | 8,329 | 8,883 | 36,788 | 0 | 0 |
| ConforMix | 2,295 | 1,590 | 23,347 | 23,217 | 3,551 |
| Boltz2 (MD) | 3,963 | 6,375 | 43,662 | 0 | 0 |
| BioEmu | 3,058 | 13,425 | 30,238 | 7,266 | 13 |

**Table S4:**
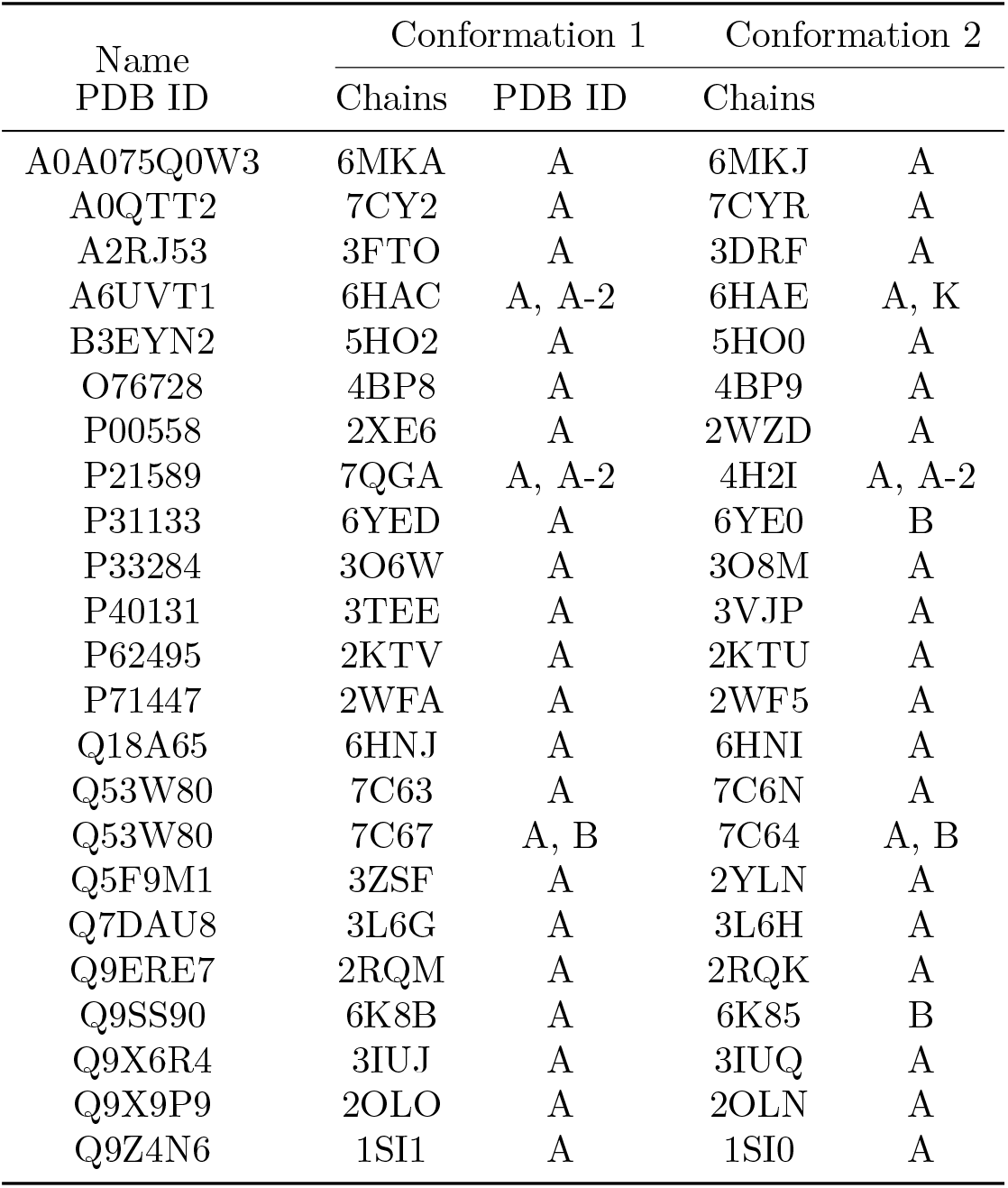
Open-closed dataset.

#### S1.4 Transporters dataset

The transporters dataset, introduced in [16] includes 15 transporter structures with inward-facing (IF) and outward-facing (OF) conformations. Although this dataset was curated from single chain structures due to the limitations of the structure prediction models at the time, for the transporters that form obligate complexes, their complete oligomeric sequences were used to perform predictions.

For the proposed IF and OF conformations of the melB transporter, both the 7L17 and 4M64 structures are classified as OF by the studies that initially reported these structures [17, 18]. Hence, the 8T60 structure with the inward-facing conformation [19] has been used with the originally proposed 7L17 outward-facing conformational state.

Similarly, the proposed IF and OF structures for the target AAC3 correspond to different proteins where the IF labeled AAC3 is the ADP/ATP translocase protein of *Thermothelomyces thermophilus* and the OF AAC3 structure is the ADP/ATP translocase 3 protein of *Saccharomyces cerevisiae*. To ensure that both conformations correspond to identical proteins, this target has been replaced with the recently experimentally determined IF and OF conformations of plastid/parasite ATP/ADP translocator [20]. The final list of targets and their PDB identifiers can be seen in Table S5.

**Table S5:** Transporters dataset.

| Name | Inward-Facing (IF) |  | Outward-Facing (OF) |  |
|---|---|---|---|---|
|  | PDB ID | Chains | PDB ID | Chains |
| A5U30_003247 | 6E9N | A | 6E9O | B |
| CYME_CMD148C | 6A6N | A, A-2 | 7DQV | A, A-2 |
| MFSD2A | 7MJS | X | 7N98 | A |
| NTT1 | 9J3N | A | 9J3O | A |
| PF0708 | 6FHZ | A | 3VVS | A |
| SLC1A1 | 6S3Q | A, B, C | 6X3E | A, B, C |
| SPF1 | 6XMS | A | 6XMU | A |
| TM_0287 | 4Q4A | A, B | 6QV1 | A, B |
| TT_C0976 | 5MKK | A, B | 6RAJ | A, B |
| mdfA | 6VS1 | A | 6GV1 | A |
| melB | 8T60 | A | 7L17 | A |
| murJ | 6NC7 | A | 6NC9 | A |
| ptsG | 6BVG | A, B | 5IWS | A, A-2 |
| slc39 | 6BTX | A | 5AYM | A |
| wlaB | 5C78 | A, D | 6HRC | A, B |

### S2 Supplementary methods

#### S2.1 Principal component analysis of RNA structures

The principal component analysis for the HIV-1 TAR and RNase P structures was performed over C1’ atom positions of nucleic acid residues of experimentally resolved structures. The AF3 and AF3 GNMCADS predicted structures are then projected onto the first two principal components calculated.

For RNase P, the cryo-em structures 9ZF2, 9ZFC, 9ZFD, 9ZGM, 9ZGN, 9ZGO, 9ZGP, 9ZGQ, 9ZGR, 9ZGS, 9ZGT, 9ZGU, 9ZGW, 9ZGV, 9ZGX, 9ZGY, 9ZGZ, 9ZH0, 9ZH1, 9ZH2, 9ZH5, 9ZH7, 9ZH9, 9ZHA, 9ZHB, 9ZHC, 9ZHD, 9ZHE, 9ZHF, 9ZHG, 9ZHI, 9ZHJ, 9ZHK, 9ZHL, 9ZHO, 9ZHP, 9ZHR, 9ZHS, 9ZHT, 9ZHU, 9ZHV, 9ZHW, 9ZHX, 9ZHZ, 9ZI2, 9ZI3, 9ZI5, 9ZF3, 9ZF5, 9ZI0, 9ZDV, 9ZI6, 9ZI4, 9ZI7, 9ZF9, 9ZI8, 9ZI9, 9ZIB, 9ZIC, 9ZID, 9ZIE, 9ZIF, 9ZIG, 9ZIH, 9ZII, 9ZIJ, 9ZIK, 9ZIL, 9ZIM, 9ZIN, 9ZIU, 9ZIV, 9ZIW, 9ZIZ, 9ZIX, 9ZIY, and 36UK [21], and the RNase P structures derived by atomic force microscopy [22] were used to perform principal component analysis.

For HIV-1, the structures 1ANR [23], 1ARJ [24], 1UTS [25], 1UUD [26], 1UUI [26], 2KDQ [27], 2KX5 [28], 2L8H [29], 5J0M [30], 5J1O [30], 5J2W [30], 6D2U [31], 7JU1 [32], 8THV [33], and 9PDG [34] were used to perform principal component analysis.

### S3 TM similarity plots

**Figure S1:**
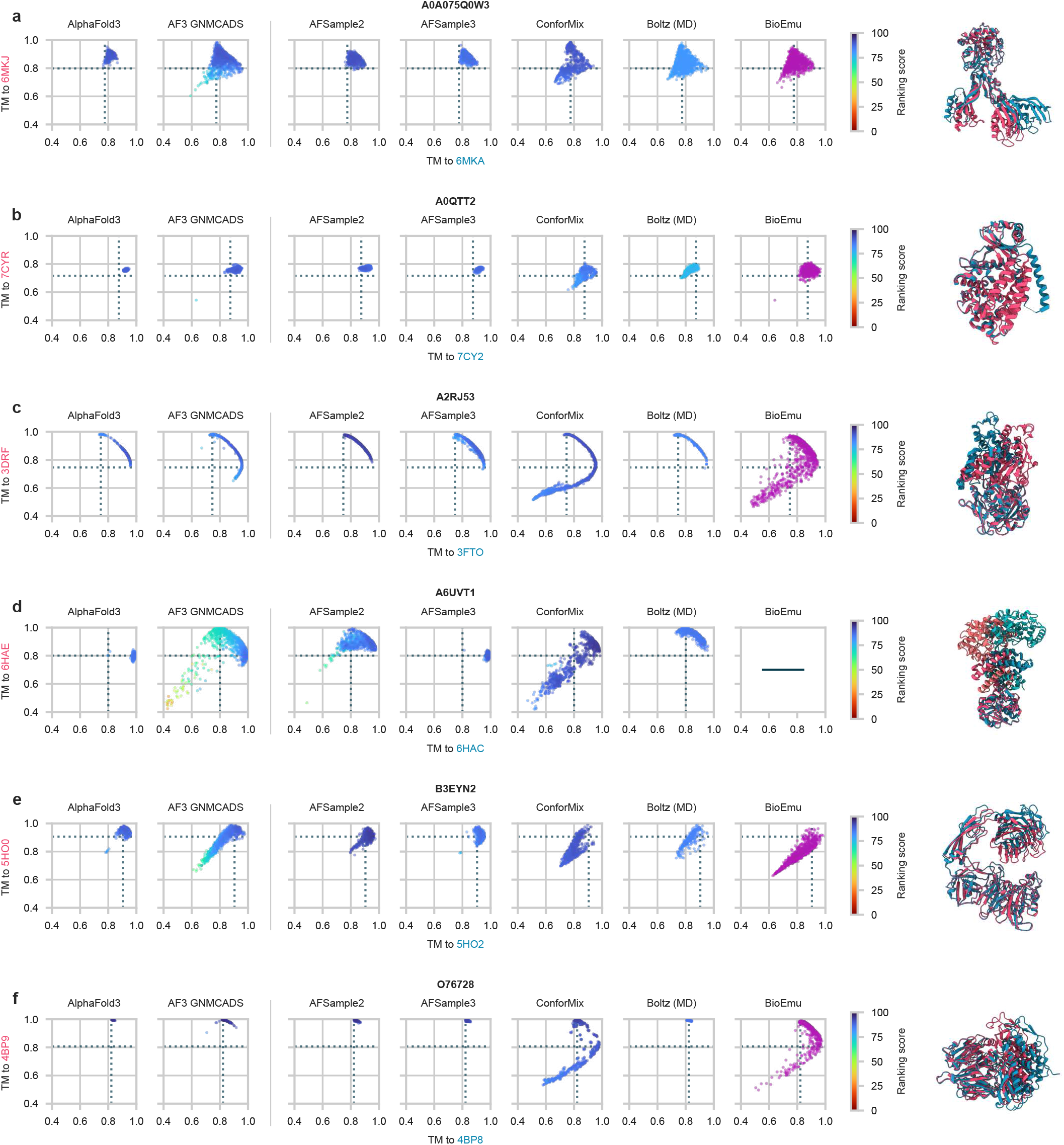
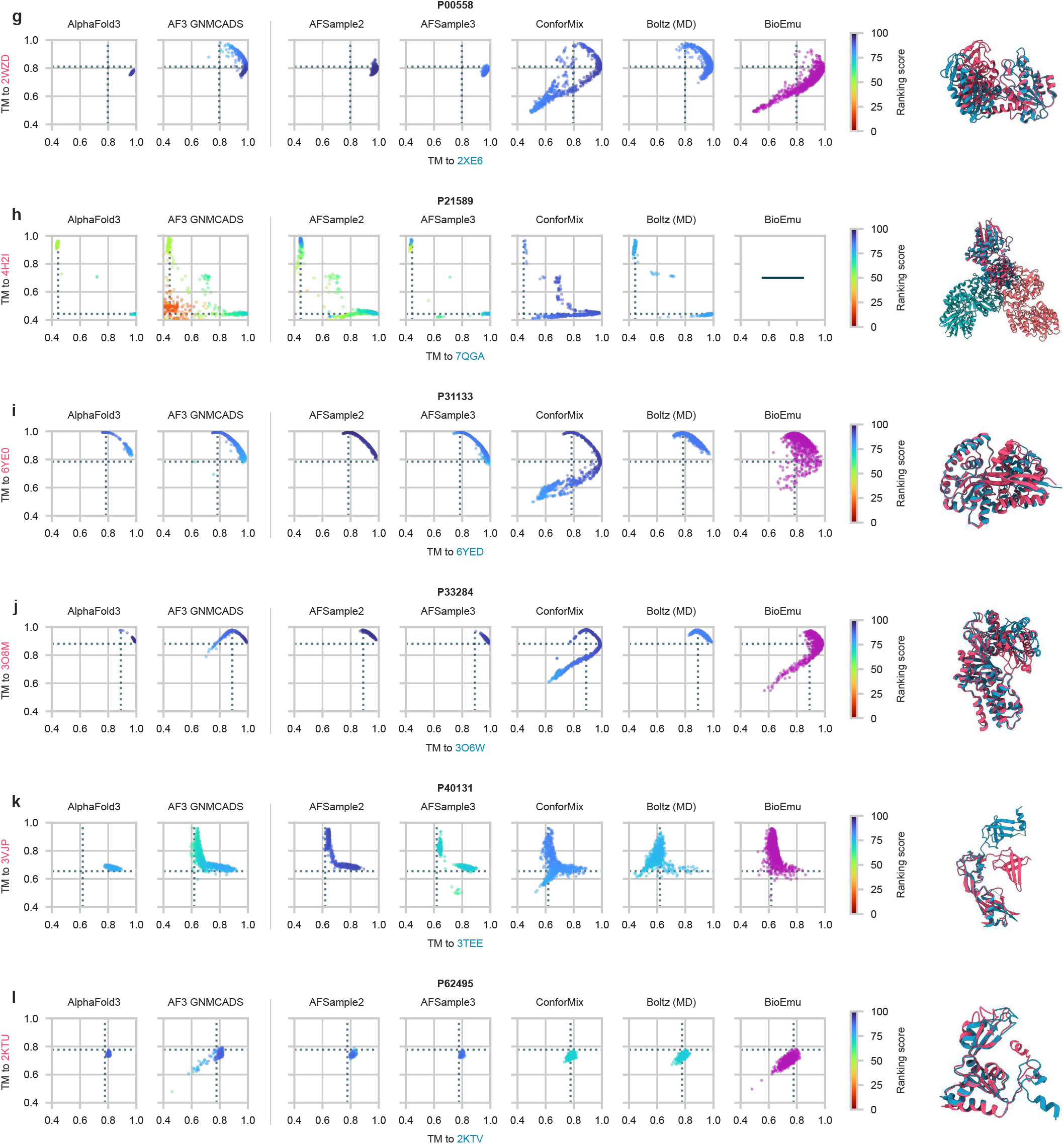
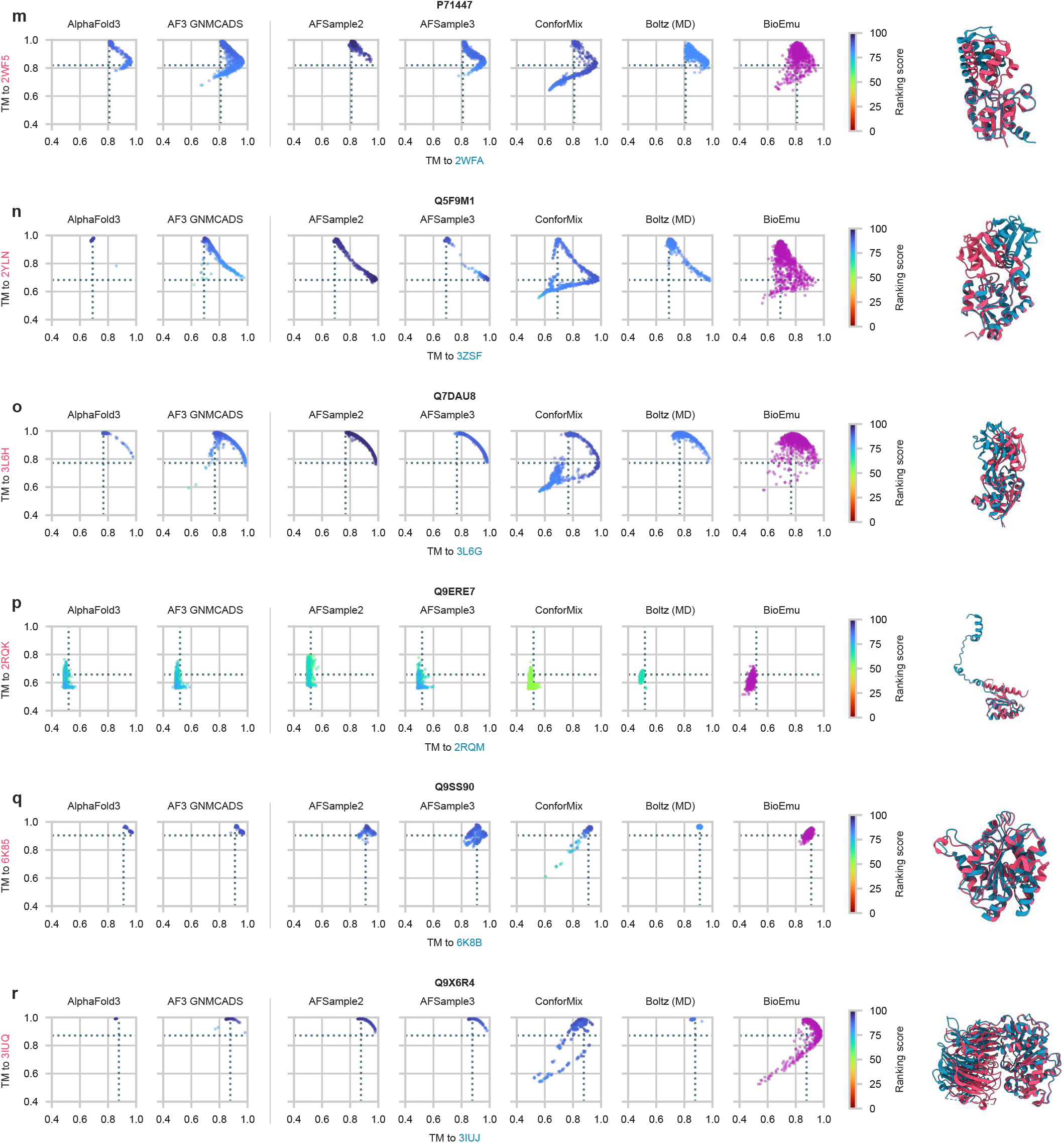
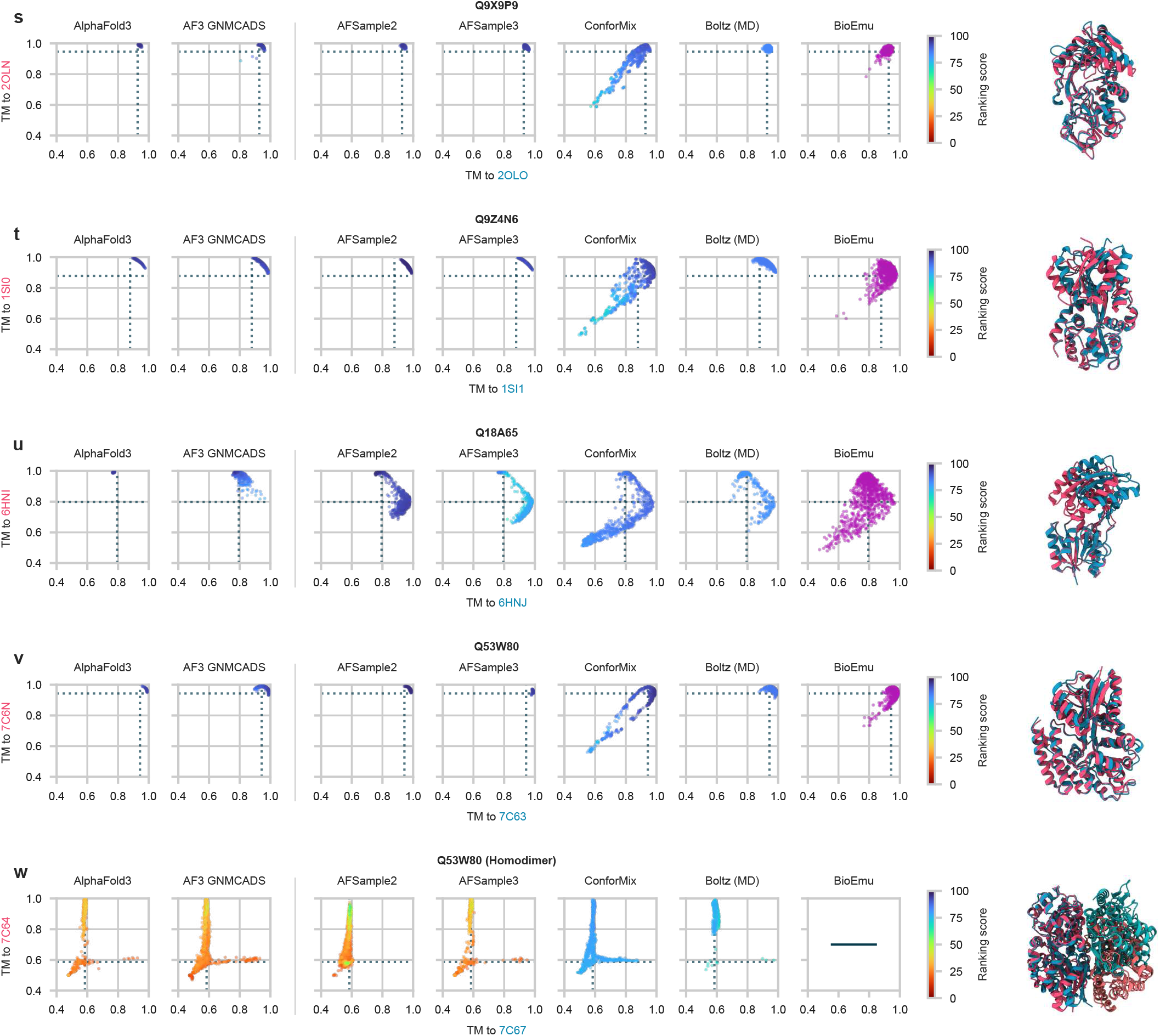
TM similarity plots for the open-closed dataset targets (continued on next page). BioEmu plot for the target A6UVT1 is not shown as BioEmu does not support multichain protein complexes. BioEmu plot for the target Q53W80 (Homodimer) is not shown as BioEmu does not support multichain protein complexes.

**Figure S2:**
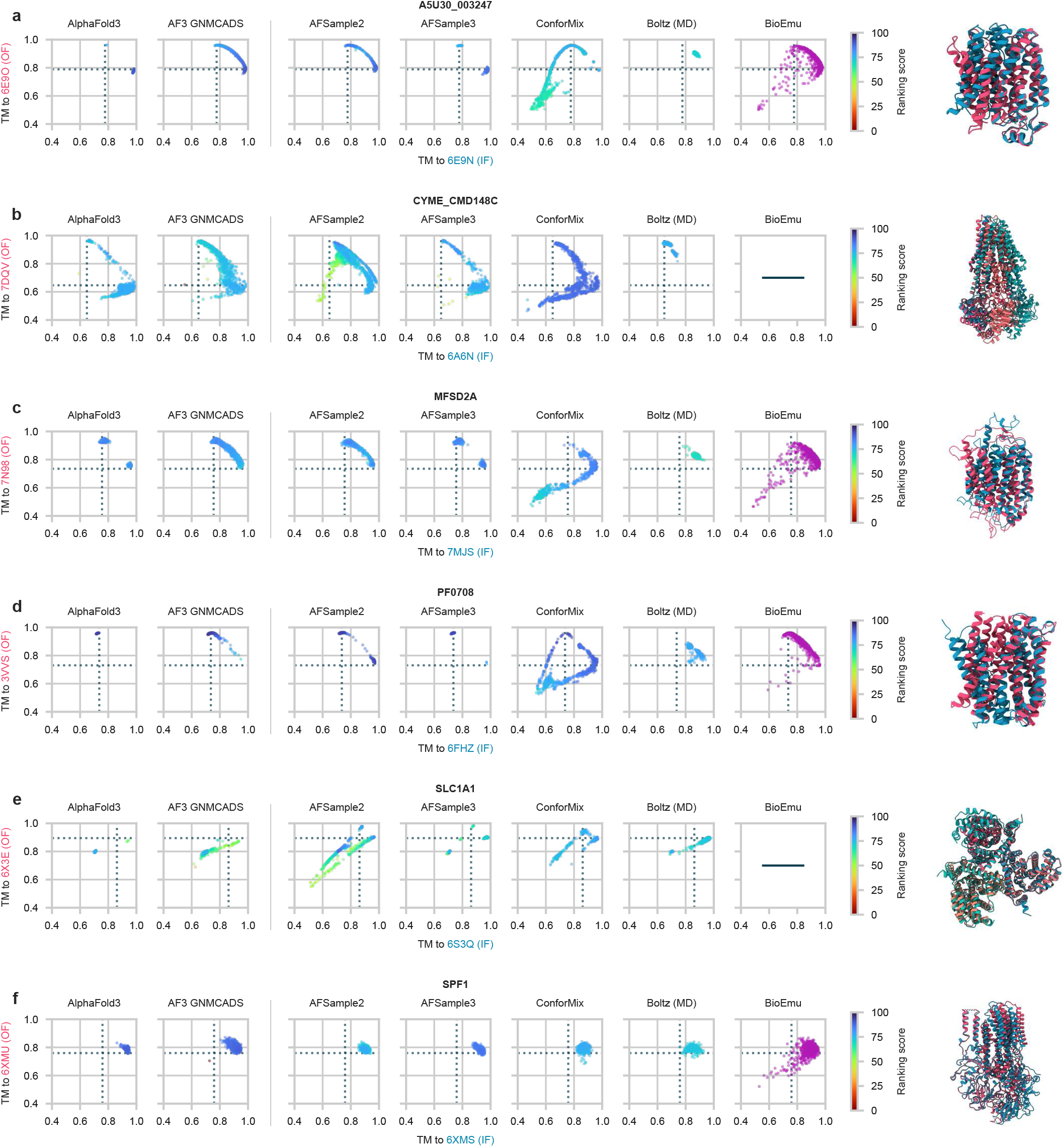
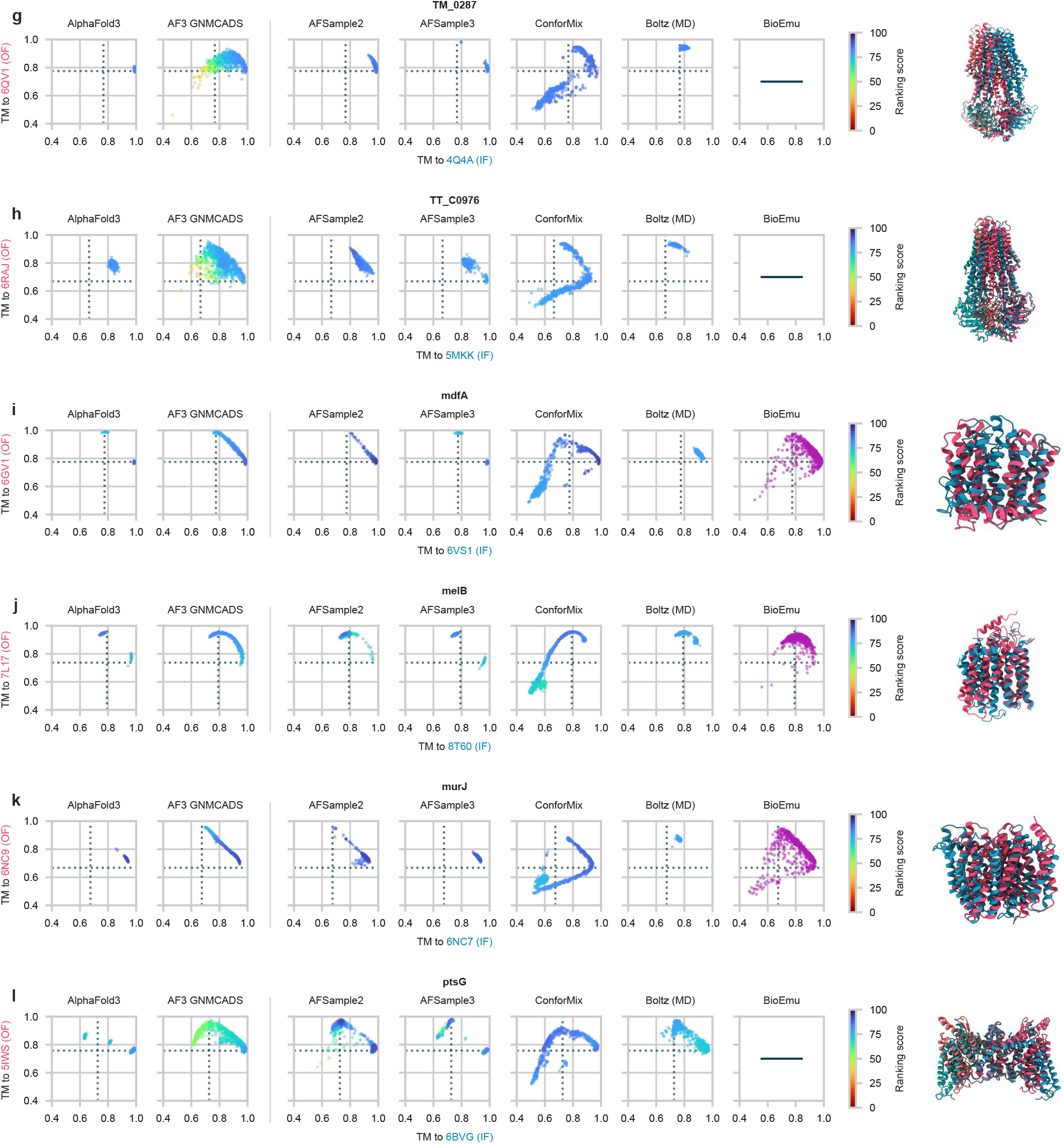
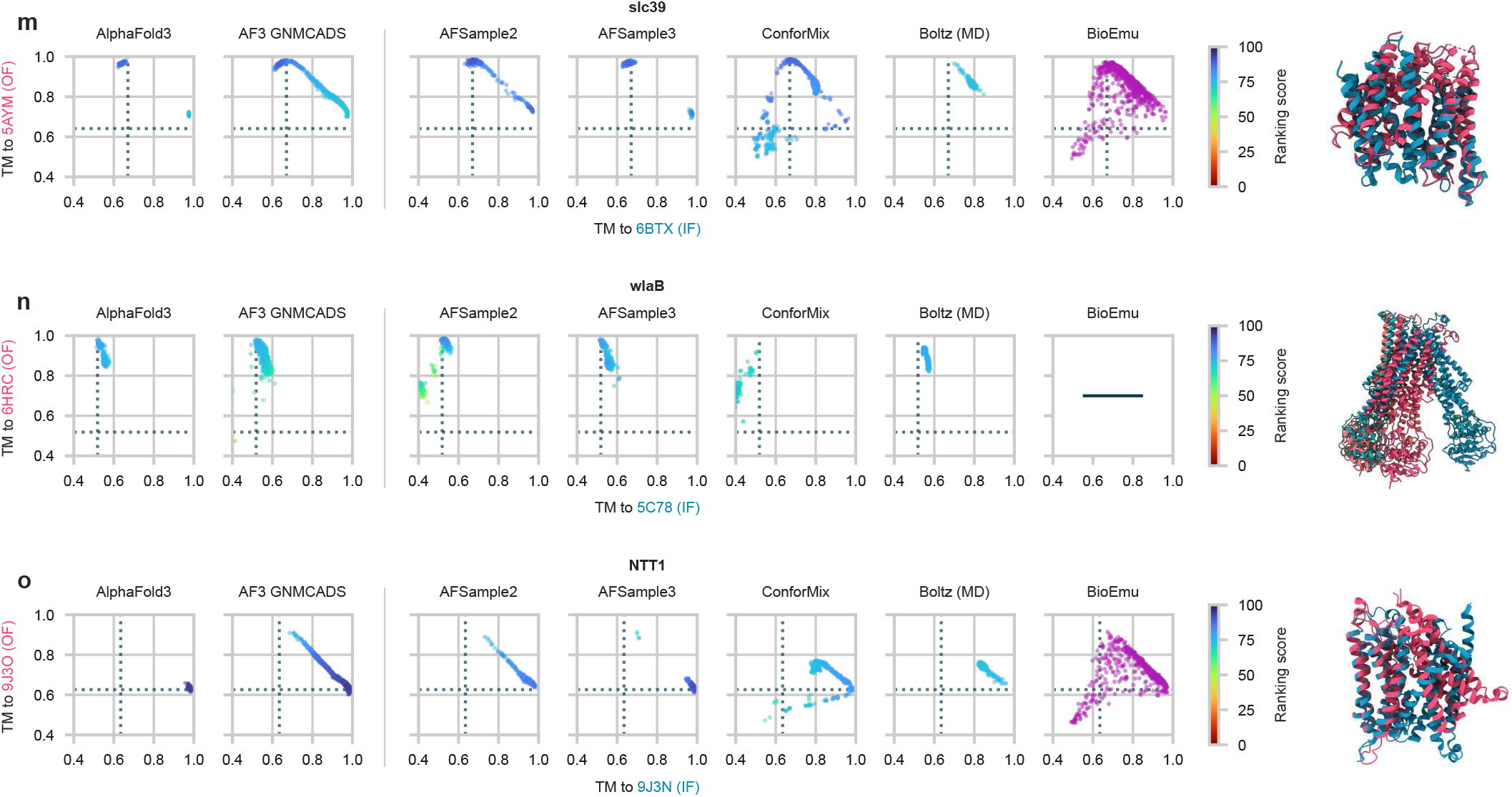
TM similarity plots for the transporter dataset targets (continued on next page). BioEmu plots for the targets CYME_CMD148C and SLC1A1 are not shown as BioEmu does not support multichain protein complexes. BioEmu plots for the targets TM_0287, TT_C0976, and ptsG are not shown as BioEmu does not support multichain protein complexes. TM similarity plots for the transporter dataset targets. BioEmu plot for the target wlaB is not shown as BioEmu does not support multichain protein complexes.

